# Modular Tissue-in-a-CUBE Platform to Model Blood-brain-Barrier (BBB) and Brain Interaction

**DOI:** 10.1101/2023.02.25.529996

**Authors:** Isabel Koh, Masaya Hagiwara

## Abstract

With the advent of increasingly sophisticated organoids, there is growing demand for technology to replicate the interactions between multiple tissues or organs. This is challenging to achieve, however, due to the varying culture conditions of the different cell types that make up each tissue. Current methods often require complicated microfluidic setups, but fragile tissue samples tend not to fare well with rough handling. Furthermore, the more complicated the human system to be replicated, the more difficult the model becomes to operate. Here, we present the development of a multi-tissue chip platform that takes advantage of the modularity and convenient handling ability of the CUBE device. We first developed a blood-brain barrier (BBB)-in-a-CUBE by layering astrocytes, pericytes, and brain microvascular endothelial cells in the CUBE, and confirmed the expression and function of important tight junction and transporter proteins in the BBB model. Then, we demonstrated the application of integrating Tissue-in-a-CUBE with a chip in simulating the testing of the permeability of a drug through the blood-brain barrier (BBB) to the brain and its effect on treating brain cancer. We anticipate that this platform can be adapted for use with organoids to build complex human systems *in vitro* by the combination of multiple simple CUBE units.

## Introduction

Neurological disorders are increasingly becoming contributors to a decreased quality of life in ageing population worldwide ^1^. Yet, progress in drug development for neurological diseases is often hampered by difficulties in getting the therapeutic substances past the blood-brain barrier (BBB) into the central nervous system. To overcome these difficulties, human *in vitro* models of the BBB are highly desired to efficiently conduct experiments to determine the permeability of a drug candidate through the BBB, as well as to study its effect on the target tissue of interest. The main components of the BBB are endothelial cells, pericytes, astrocytes, and a basement membrane that together contribute to maintaining the barrier function and homeostasis of the brain via tight junction and transporter proteins ^2–4^. Given the complex makeup of the BBB, various models have been established with different combinations of cellular and basement membrane components – astrocytes, pericytes and brain endothelial cells from primary, immortalized, or pluripotent stem cell (PSC)-derived sources have been used with synthetic membranes or hydrogels as basement membrane ^5–14^.

Besides the cellular and structural components of the BBB, the interaction between the BBB and the brain is also important to be taken into consideration in developing BBB models. In neurodegenerative diseases and in the ageing brain, a cycle of the accumulation of pathological proteins such as amyloid-β in Alzheimer’s disease or Lewy bodies in Parkinson’s disease leading to dysfunction of the BBB, leading to further accumulation of pathological proteins and further disruption of the BBB, is thought to contribute to the progression of the disease ^15–18^. Thus, to effectively develop and test drug candidates to treat these diseases requires recapitulating not just the healthy BBB, but also the dynamics of the BBB-brain interaction, particularly in the diseased state.

Organoids derived from stem cells have been shown to mimic some of the function of native organs, and with the rapid expansion of disease-modelling brain organoids being developed ^19,20^, there is increasing demand to progress *in vitro* BBB models to include a brain component to more closely represent *in vivo* conditions. Combining different tissue models to achieve a more physiological representation of an organ, commonly termed Organ-on-a-Chip systems, can be achieved in two main ways: (i) by simple merging of the separately cultured tissues such as co-culturing retinal organoid on a bed of retinal pigment epithelium (RPE) cells ^21^ or brain organoid on a vascular network ^22^, or (ii) by connecting organoids cultured in modular components via microfluidic channels, for example in a gut-liver model ^23^ or a combination of 4 to 6 multi-organ model ^24,25^.

Lamentably, many of these engineered models are not widely adopted by biology-based laboratories that study disease mechanisms or drug discovery and development, ostensibly because they do not have the time or resources for the setup processes that can be long and complicated for those unfamiliar with the technologies. Furthermore, in most Organ-on-a-Chip systems, the cells adhere and grow on the device, making it difficult to retrieve the sample for later use or analysis without detaching the cells from the device, which can cause damage to the sample. Hence, there is a need to balance replicating *in vivo* complexities in the lab, but with simple setup procedures that can be handled by researchers without an engineering background.

In this paper, we present a highly modular and adaptable platform for Organ-on-a-Chip development, which takes advantage of the ease of sample handling of our previously published CUBE culture device ^26–28^ to effortlessly integrate multiple separately cultured tissue or organoid models in a simple chip device (Fig. 1). The desired tissue or organoid can be reconstructed in the CUBE with a combination of the appropriate cell types and extracellular matrix (ECM) hydrogel that provides a 3D scaffold for cells. As the cells are cultured in modular units, the timing to start the experiments can be controlled by accounting for different maturation times of different tissues or organoids. Additionally, the chip device can be disassembled at the end of the experiment to retrieve the sample for further experiments or analyses.

**Figure 1.**
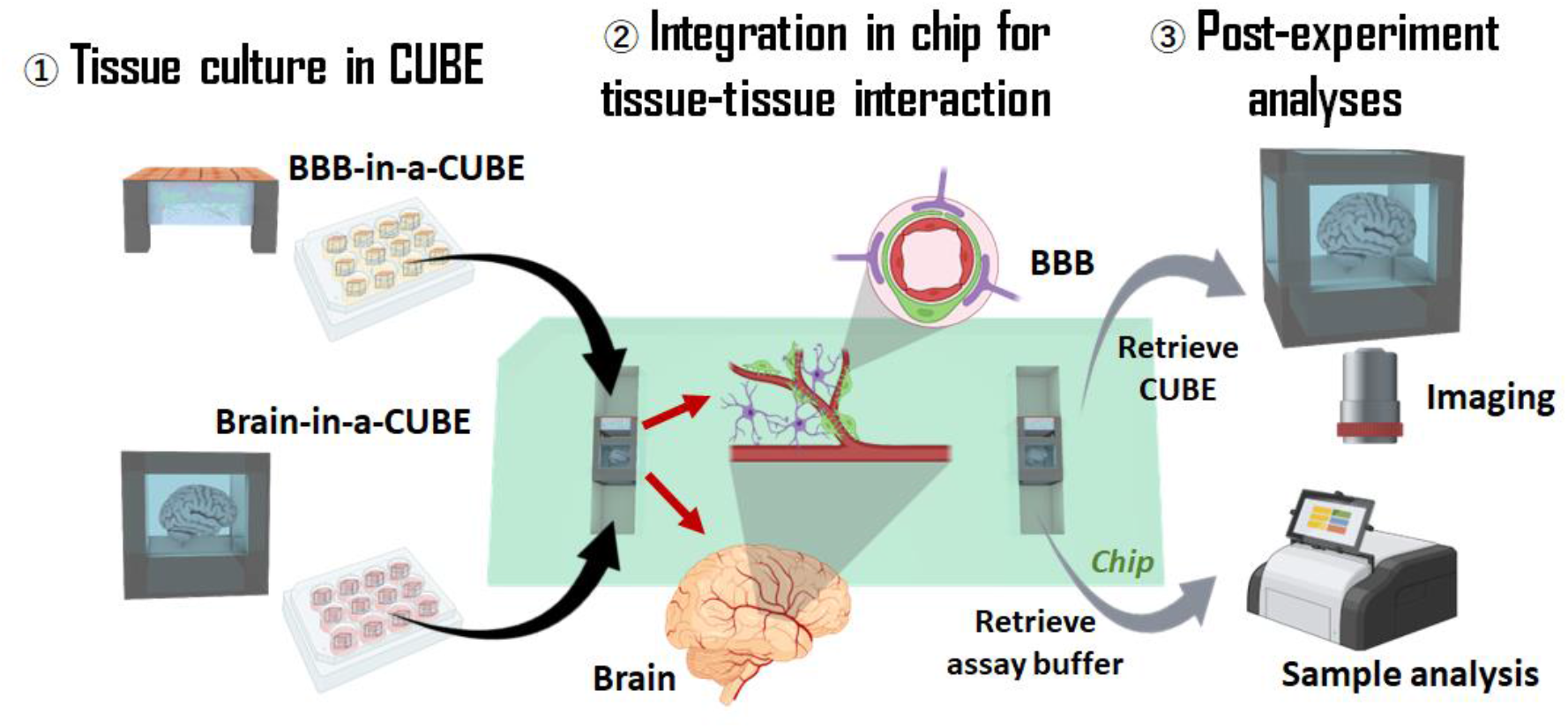
Integration of multiple Tissue-in-a-CUBEs for multi-tissue interaction. The concept of this work was to utilise the CUBE culture device to facilitate the modular combination of complex 3D tissues in simple units to build complex systems. (1) Modular tissue units are first cultured in individual CUBEs, (2) then transferred to a chip at the appropriate timing to initiate tissue-tissue interaction, and (3) at the end of the experiment, samples can be retrieved from the chip for analyses.

By employing this platform, we demonstrate the application of this Organ-on-a-Chip platform with a BBB-brain model. We first developed a BBB-in-a-CUBE model by co-culturing primary astrocytes and pericytes in a basement membrane hydrogel (Matrigel) with iPSC-derived BMECs seeded on the top surface of the Matrigel, and confirmed the expression of important tight junction and transporter proteins by immunofluorescence staining and RT-qPCR. We also validated the function of the BBB by measuring the trans-endothelial electrical resistance (TEER), as well as conducting permeability experiments with Lucifer Yellow (passive paracellular diffusion) and Rhodamine123 (P-glycoprotein, PGP substrate transport). Finally, as a proof-of-concept, we designed a BBB-brain chip to contain a BBB-in-a-CUBE and a Glioblastoma-in-a-CUBE model to demonstrate, as reported by Tivnan *et al*. ^29^, that the BBB blocking of the PGP substrate drug Vincristine into the brain can be overcome with the addition of PGP inhibitor Reversan. Thus, we show that this platform can be utilized to achieve complex systems by the combination of several small and simple organoid units, without the need for complicated external pump or power sources.

## Materials and Methods

### CUBE and Chip Fabrication

CUBEs, moulds to fabricate the chip, and chip base holders were designed using Rhinoceros 3D software (Robert McNeel & Associates) and ordered from a machining company (Proto Labs Japan). All CUBEs, chip moulds, and chip base holders were made of aluminium. Design dimensions and methods to fabricate PDMS are shown in supplementary figure S1.

The original CUBE design was modified with a thicker frame on the top side of the CUBE to allow the attachment of a nitrile O-ring (AS ONE, 62-3049-63) to the CUBE, which was necessary to ensure a tight seal between CUBE and chip ^30^. Two types of CUBEs were used in this study: (i) for BBB-in-a-CUBE, the bottom half of the CUBE was removed to reduce the amount of Matrigel used, and (ii) for Glioblastoma-in-a-CUBE, the sidewalls of the CUBE were covered with PDMS (Fig. S1A). PDMS (Silpot 184, Dow Toray, 04133124) was prepared by mixing the elastomer base with the curing reagent at a 10:1 ratio. To make PDMS sidewall with a thickness equivalent to that of the CUBE frame (0.75 mm), ∼2.8 g of the mixture was spread out in a 100 mm dish and degassed to remove air bubbles, before placing CUBE frames on the dish. The PDMS was degassed again and baked at 85°C for 20∼30 min. After curing, excess PDMS was trimmed from the frames with a scalpel, and the process repeated for the remaining three adjacent sides of the CUBE, leaving the top and bottom surfaces open (Fig. S1B).

Two types of chips were used in this study: (i) for TEER measurements and permeability tests, the chip moulds were designed to hold the BBB-in-a-CUBE with two media chambers at the top and bottom sides of the CUBE, and (ii) for BBB-brain interaction experiments, the chip moulds were designed to fit BBB-in-a-CUBE and Glioblastoma-in-a-CUBE together, with media chambers on the two ends of both CUBEs (Fig. S1C). Each chip comprises a base component which holds the CUBEs and media compartments, and a lid component with access ports to seal the chip. PDMS chips were made by pouring uncured PDMS into the mould, degassing to remove air bubbles, then baked at 85°C for about 1 hr. After curing, the PDMS chips were pried out of the mould (Fig. S1D).

Prior to use with cell culture, CUBEs and chips were washed with ultrasonication once in MilliQ water and twice in isopropanol (IPA), for 10 min each wash, then dried in the oven for 2 hr. O-ring was washed as follows: soak in acetone for ∼4 hours, discard acetone and replace with fresh acetone to soak overnight, discard acetone and ultrasonicate with IPA for 10 min, discard IPA and ultrasonicate with autoclaved MilliQ water for 10 min, then spread O-rings out in a dish and leave to dry overnight. To assemble the CUBE and chip setup, a double-sided adhesive film (NSD-100, NIPPA) was attached onto the lid component of the chip and holes punched in the access ports to allow media to be added to the chip later. The chip base was placed in a holder to prevent it from overexpanding when it is sealed. Then, the CUBEs were placed in the base component, and the adhesive lid placed on top of the base to seal the chip. The sealed chip was then placed in a clamp holder to ensure a tight seal of the PDMS chip, and media can be added to the media chambers via the access ports. The clamp holder was purchased from Micronit (Fluidic Connect PRO Chipholder Frame; FCPROCH) with an attachment customized to fit our chip design ordered from a microfabrication company (Icomes Lab, Japan) (Fig. S1E, supplementary video SV1).

### Cell culture

Primary normal human astrocytes (NHA; Lonza, CC2565) were cultured in astrocyte growth medium (AGM Bulletkit; Lonza, CC3186) supplemented with 0.7% Penicilin-Streptomycin (PS; Gibco, 15140122). Primary human brain vascular pericytes (HBVP; ScienCell, 1200) and glioblastoma multiforme cell line T98G (RIKEN Cell Bank, RCB1954) were cultured in DMEM/F12 with GlutaMAX supplement (Gibco, 10565018) supplemented with 10% foetal bovine serum (FBS; Cytiva, SH30396.03) and 1% PS. NHA, HBVP, and T98G were dissociated using 0.25% Trypsin-EDTA (Gibco, 25200-056) and trypsin neutralizing solution (TNS; Kurabo, HK3220). NHA and HBVP were used within passages p4∼7, and T98G used within p7∼13 in experiments. IMR90-4 iPSCs (WiCell, WB65317) between passages p35∼45 were maintained on Matrigel-coated dishes in mTeSR Plus medium (STEMCELL Technologies, 100-0276) and dissociated using ReLeSR (STEMCELL Technologies, 05872). For Matrigel coating, hESC-qualified Matrigel (Corning, 356231) was diluted 1:100 in DMEM/F12 according, and 1 mL of the diluted Matrigel was used to coat one 35 mm culture dish. The dish was incubated at room temperature for one hour, and rinsed once with 1x DPBS (Gibco, 14190) before use. For use in differentiation, IMR90-4 were dissociated using Accutase (Invitrogen, 00-4555-56) and plated with mTeSR Plus with 10 μM Rock inhibitor (Y27632; Nacalai Tesque, 08945-84) for the first 24 hr, then without Y27632 from the next day. When the IMR90-4 have reached ∼70% confluency, differentiated to BMEC was initiated according to the protocol of Lippmann *et al*. ^7,31^. On Day 0 of differentiation, medium was switched to neural-endothelial differentiation medium which was DMEM/F12 Ham without L-glutamine (Sigma-Aldrich, D6421) supplemented with 20% KnockOut serum replacement (KOSR; Life Technologies, 10828010), 1% MEM non-essential amino acid (NEAA; Nacalai Tesque, 06344-56), and 1% GlutaMAX supplement (Gibco, 35050061). After 6 days of neural-endothelial differentiation, the medium was switched to BMEC maturation medium, which comprises human endothelial serum-free medium (hESFM; Gibco, 11111044) with 1% human platelet poor-derived serum (HS; Sigma-Aldrich, P2918), 20 ng/mL basic FGF (bFGF; Fujifilm Wako, 060-04543), and 10 μM all-trans retinoic acid (ATRA; Nacalai Tesque, 36331-44). To coat a dish with collagen IV (COL IV) and fibronectin (FN), 4 mL of a mixture of 50 μg/mL COL IV (Sigma-Aldrich, C7521) and 25 μg/mL FN (Sigma-Aldrich; F2006) in DPBS was used to coat one 100 mm culture dish and incubated at 37 °C for at least 2 hr. The excess solution was aspirated and left to dry overnight before use. After 2 days of BMEC maturation, cells were dissociated using Accutase for 20-25 min and re-plated on COLIV+FN-coated dishes in BMEC maturation medium and incubated for 1 hr. After 1 hr, the cells were washed twice gently with 1x DPBS, then fresh BMEC maturation medium was added, and the cells incubated for another day before use in making BBB. To image the 3D structure of BBB-in-a-CUBE, astrocytes were labelled with CellLight Actin-GFP (Invitrogen, C10582), pericytes labelled with CellLight Tubulin-RFP (Invitrogen, C10614), and BMECs labelled with CellLight Plasma membrane-CFP (Invitrogen, C10606) overnight, according to the manufacturer’s protocol, prior to making BBB-in-a-CUBE.

### BBB-in-a-CUBE

Before seeding cells in the CUBE, an O-ring was attached to the thicker part of the CUBE frame. Astrocytes (0.5×10^6^ cells/mL) and pericytes (0.5×10^6^ cells/mL) were suspended in Matrigel, and 20 μL of the cell suspension was added to each CUBE, before incubating at 37 °C for 25 min for Matrigel to cure. After Matrigel has cured, 10 μL of BMEC (5.5×10^6^ cells/mL suspended in BMEC maturation medium with 10 μM Y27632) was seeded on the top surface of the gel, then incubated at 37 °C for 1 hr for BMEC to attach onto the Matrigel. After BMECs have adhered, the CUBEs were transferred to a 48-well plate containing BMEC maturation medium with 10 μM Y27632. The following day, medium was switched to BBB medium which comprises a 1:1 mix of EC medium (hESFM+1% HS) and AGM. Medium was changed every other day (on Days 3 and 5) by discarding and replacing with fresh half the volume of medium. BBB-in-a-CUBE were used at Days 2, 4, and 6 for TEER measurements and Lucifer Yellow permeability tests, and at Day 6 for Rhodamine123 permeability tests and BBB-brain experiments. A CUBE with 20 μL Matrigel only without cells was used as a Blank or No Cell condition.

### Glioblastoma-in-a-CUBE

Before seeding cells in the CUBE, an O-ring was attached to the thicker part of the CUBE frame. T98G cells were suspended in Matrigel at 1.0×10^6^ cells/mL, then added to the CUBE and incubated at 37 °C for 25 min for Matrigel to cure. After curing, the CUBEs were transferred to a 48-well plate containing T98G medium. Glioblastoma-in-a-CUBE were used at Day 3 for BBB-brain experiments.

### Immunofluorescence Staining

To fix the cells, BBB-in-a-CUBE samples were washed with DPBS twice, fixed with 4% paraformaldehyde for 20 min, then washed with DPBS for 10 min twice. Samples for cryo sectioning were soaked successively in 10%, 15%, and 20% sucrose (Wako, 194-00011) for 1 hr each, then in cryosection embedding medium (Tissue-Tek O.C.T compound; Sakura Finetek, 4583) for 10 min. After this, samples were removed from the CUBE and placed in a mould filled with OCT, then frpzen at -80°C for 30 min. Once frozen, samples were removed from the mould and sectioned at 50 μm thickness using a microtome equipped with a freezing unit (Yamato Kohki Industrial, Retoratome REM-710 and Electro Freeze MC-802A). Sections were collected on glass slides and dried with cool air for 30 min, then in a 40 °C oven for another 30 min, followed by soaking in MilliQ water to remove excess OCT. After removing excess water, the sections were dried at RT for 10 min.

For immunofluorescence staining, permeabilization was performed by incubating samples in 0.5% Triton X-100 for 10 min, then washing with 100 mM Glycine for 10 min three times. Immunofluorescence buffer (IF buffer) consisted of 0.5% Tween20, 2% Triton X-100, and 10% bovine serum albumin (BSA; Sigma, 126615) in DPBS. Samples were blocked in IF buffer with 10% goat serum (Gibco, 16210064) (IF+G) for 30 min, followed by IF+G with 1% goat anti-mouse IgG (Bethyl Laboratories, A90-116A) for 20 min. Antibodies were prepared according to the dilutions in Table 1. Primary antibody incubation was overnight at 4 °C and secondary antibody incubation was 2 hr at RT. After each antibody incubation, samples were washed with IF buffer for 15 min three times. Nuclei were stained with DAPI for 20 min, then washed with DPBS for 5 min three times. Samples were then immersed in RapiClear clearing solution (SUNJin Lab, RC147001) overnight. Imaging was performed using a confocal microscope (Leica Microsystems, TCS SP8 Lightning).

**Table 1.** List of antibodies for immunofluorescence staining.

| **Reagents** | **Source, Identifier** | **Dilution** |
| --- | --- | --- |
| Anti-SLC7A5/LAT1 Rabbit monoclonal | Abcam, ab208776 | 1:200 in IF |
| Anti-MRP1 Rabbit monoclonal | Abcam, ab23383 | 1:100 in IF |
| Anti-GLUT1 Rabbit monoclonal | Abcam, ab115730 | 1:100 in IF |
| Anti-BCRP/ABCG2 Rabbit monoclonal | Abcam, ab207732 | 1:100 in IF |
| Anti-PGP/MDR1 Rabbit polyclonal | Abcam, ab235954 | 1:100 in IF |
| Anti-OAT3 Rabbit polyclonal | Abcam, ab247055 | 1:100 in IF |
| Anti-Serotonin Rabbit polyclonal | Abcam, ab272912 | 1:500 in IF |
| Anti-MCT1 Rabbit polyclonal | Abcam, ab85021 | 1:20 in IF |
| Anti-ZO1 Rabbit polyclonal | Abcam, ab216880 | 1:200 in IF |
| Anti-Claudin5 Rabbit polyclonal | Abcam, ab15106 | 1:200 in IF |
| Anti-CD31 Mouse monoclonal | Dako, M0823 | 1:50 in IF |
| Anti-von Willebrand Factor Sheep polyclonal | Abcam, ab111713 | 1:50 in IF |
| Goat anti-rabbit AF488 | Invitrogen, A32731 | 1:200 in IF |
| Goat anti-rabbit AF647 | Invitrogen, A32733 | 1:200 in IF |
| Goat anti-mouse AF555 | Invitrogen, A32727 | 1:200 in IF |
| Dog anti-sheep AF555 | Invitrogen, A21436 | 1:200 in IF |

### Reverse Transcription Quantitative Polymerase Chain Reaction (RT-qPCR)

Cells were harvested from BBB-in-a-CUBE by detaching the BBB from the CUBE into cold DPBS in a dish, then transferring the BBB to a tube with Cell Recovery Solution (Corning, 354253) for 20 min on ice. Cells were pelleted by centrifugation, and washed with cold DPBS after discarding the supernatant. Centrifugation and DPBS wash steps were repeated before proceeding with RNA extraction. Total RNA was extracted using Total RNA Extraction Miniprep System (Viogene, GR1001) and quantified using Eppendorf BioSpectrometer basic. 4∼13 samples from one biological replicate were pooled together in one tube, and the volume of total RNA equivalent to one sample was used to prepare the cDNA using SuperScript IV VILO Master Mix with ezDNase Enzyme (Invitrogen, 11766050) according to the manufacturers’ protocols; 3 biological replicates were used in this study. qPCR was carried out using SYBR Green Realtime PCR Master Mix (Toyobo, QPK-201) according to the manufacturer’s protocol for use with amplification and detection by Analytik Jena qTower^3^: 95 °C for 1 min, and 40 cycles of 95 °C for 15 s, 60 °C for 30 s, and 72 °C for 1 min. Primer sequences are listed in Table 2. Primers were ordered from Eurofins Scientific with sequences for ZO-1, CLDN5, OCLN, JAM-A, MDR1-^*^2, and MDR1-^*^3 obtained from OriGene Technologies, and sequences for MDR1-^*^1, MRP1, BCRP, GLUT1, LAT1, MCT1, OAT3, and AQP4 obtained from Kurosawa *et al*. 2018 ^32^. The mRNA expression level was calculated as the fold change 2^−ΔCt^, where ΔC_t_ is obtained by subtracting Ct of TATA-binding protein from the C_t_ of the target gene.

**Table 2.**
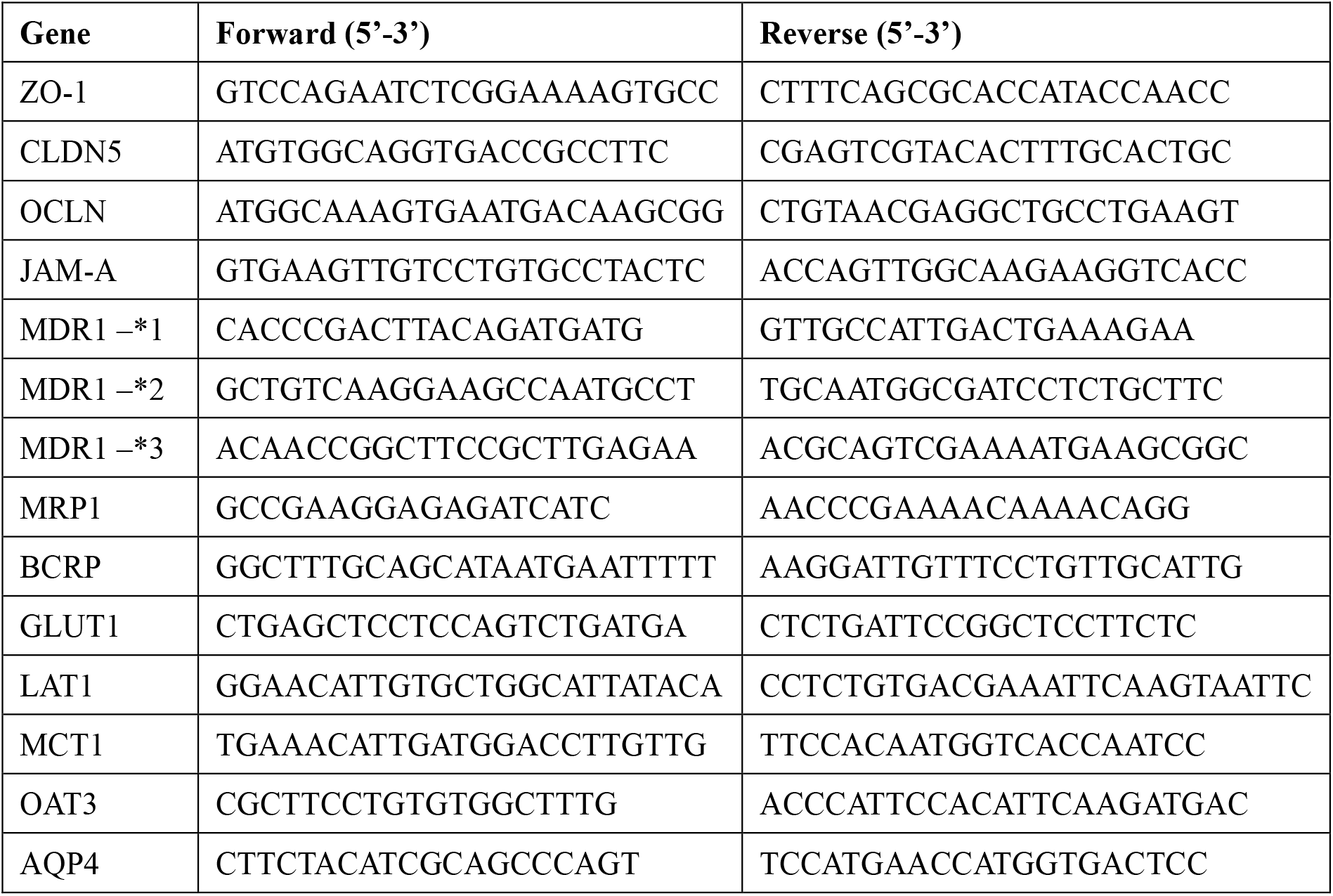
Forward and reverse primer sequences for qPCR.

### Trans-endothelial electrical resistance (TEER) Measurement

Assay buffer comprised 0.01 M HEPES (Gibco, 15630-080), 1 mM sodium pyruvate (Nacalai Tesque, 06977-34), and 0.5% dimethyl sulfoxide (DMSO; Nacalai Tesque, 13445-74) in HBSS buffer (Nacalai Tesque, 09735-75), pH 7.4. To measure TEER, CUBEs were transferred into the TEER chip and assembled as described above. 200 μL of assay buffer warmed to 38 °C was added to each medium chamber and electrical resistance, R was measured using Millicell ERS-2 Volt/Ohmmeter (Merck, MERS00002). All procedures were performed on a 38 °C hotplate. TEER was calculated by subtracting the resistance of the BBB sample (R_BBB_) by that of the blank sample (R_blank_) and multiplying by the surface area of the gel (SA = 0.1225 cm^2^). TEER was measured for Days 2, 4, and 6, and the samples were used for Lucifer Yellow permeability tests immediately after TEER measurements. 6 samples were measured for each biological replicate, and 3 biological replicates were used in this study.

### Permeability Test

The setup for permeability tests is the same as for TEER as described above. 200 μL of assay buffer warmed to 38 °C was added to each medium chamber and the samples incubated at 37 °C for 30 min. To start the permeability test, the assay buffer was discarded, and fresh assay buffer added to the basal (astrocyte/pericyte) side of the CUBE while assay buffer with 10 μM Lucifer Yellow (LY; Fujifilm Wako, 125-06281) or Rhodamine123 (Rho123; Fujifilm Wako, 187-01703) was added to the apical (BMEC) side. For Rho123 experiments, samples were incubated with 40 μM PGP inhibitor Reversan (Sigma-Aldrich, SML0173) or an equal volume of DMSO as control for 4 hr prior to permeability experiments. At the designated time points (15, 30, 60, and 120 min), 50 μL of assay buffer was collected from the basal side for fluorometric measurement and replaced with the same volume of fresh buffer. A microplate reader (Molecular Devices, SpectraMax iD3) was used to measure the concentration of LY (Ex. 488 nm; Em. 575 nm) and Rho123 (Ex. 428 nm; Em. 536 nm). Apparent permeability, P_app_ was calculated as follows:

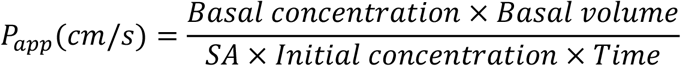

6 samples were measured for each biological replicate, and 3 biological replicates were used in this study. Samples with abnormally high concentration at *t* = 15 min (greater than 0.01 μM) were considered to have suffered damage or there was leakage in the chip, and the results were discarded.

### Tissue-tissue Interaction

For BBB-Glioblastoma experiments, BBB-in-a-CUBE at Day 6 were incubated with 40 μM Reversan or an equal volume of DMSO as control for 4 hr before being transferred to a chip with Glioblastoma-in-a-CUBE at Day 3, and the chip was assembled as described above. Glioblastoma medium was added to the basal side and BBB medium with 5 μM Vincristine (Tocris, 1257) added to the apical side. Following a 4 hr incubation at 37 °C, the media were discarded and Glioblastoma CUBEs retrieved from the chip. The CUBEs were washed once in live cell imaging solution (LCIS; Invitrogen, A14291DJ), then incubated with 1.5 μM Calcein-AM (Nacalai Tesque, 19177-14) and 1.5 μM propidium iodide (Nacalai Tesque, 19174-31) diluted in LCIS for 20 min at 37 °C. Samples were washed once in LCIS before LIVE/DEAD imaging with a fluorescence microscope (BZX-700, Keyence). The number of live and dead cells were counted using Imaris software (Bitplane, v9.0.2). 4∼5 samples were measured for each biological replicate, and 3 biological replicates were used in this study.

## Results and Discussion

The design concept for the organ-on-a-chip platform of this study was to reconstruct separately functional tissue units of an organ in a CUBE culture device, before integrating them together in a chip device to replicate complex tissue-tissue interactions. To demonstrate this concept, we first developed a BBB-in-a-CUBE model and confirmed its barrier and transport functions. Following this, we integrated the BBB-in-a-CUBE with a Glioblastoma-in-a-CUBE to convey the potential application of the platform in drug testing research.

### Reconstruction of BBB in a CUBE

The function and homeostasis of the barrier between the blood and the brain is regulated by the synergistic interaction between brain microvascular endothelial cells (BMECs) with the astrocytes and pericytes that surround them ^4,33–38^, and most previously developed *in vitro* BBB models include one or more of these three cell types in their models with or without ECM ^5–14,39^. In our model, we reconstruct the 3D structure of the BBB by first embedding primary human astrocytes and pericytes in Matrigel in the CUBE, followed by seeding hiPSC-derived BMECs on the top surface of the Matrigel (Fig. 2A) to replicate the physiological architecture of the BBB and the interactions between astrocytes, pericytes, and BMECs. BMECs tend to form tubular network structures instead of monolayers when seeded on Matrigel ^40^. However, we found that the addition of the selective Rho-associated protein kinase (ROCK) inhibitor Y-27632, which has been shown to enhance endothelial cell adhesion to substrates and promote wound healing ^41,42^, enabled the seeding of BMECs as a monolayer on Matrigel. Imaging of the fluorescently-labelled astrocytes, pericytes, and BMECs showed that over the course of 6 days of culture, astrocytes and pericytes that were initially rounded in shape at Day 2 after seeding elongated and populated the area under the BMEC layer, indicating self-organisation of the cells to form the BBB structure in 3D (Fig. 2B, supplementary figure Fig. S2∼4).

**Figure 2.**
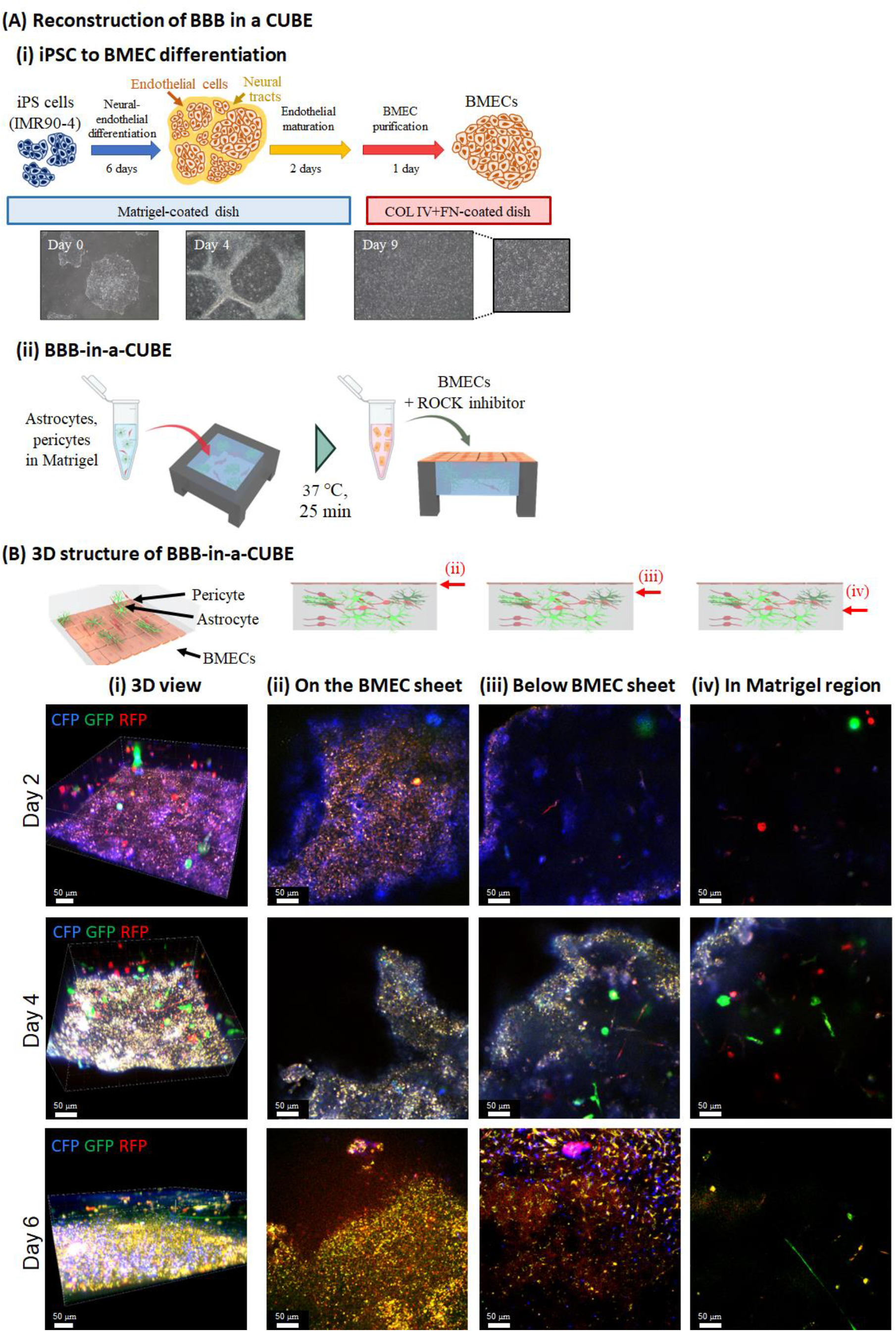
Reconstruction of the blood-brain barrier (BBB) in a CUBE. (A) The structure of the BBB was reconstructed in a CUBE using primary astrocytes and pericytes, and iPSC-derived brain microvascular endothelial cells (BMECs). (i) Differentiation of iPSC to BMEC was based on the protocol by Lippmann et al. (ii) BBB was assembled in the CUBE by first seeding astrocytes and pericytes embedded in Matrigel in the CUBE, then seeding BMECs with ROCK inhibitor Y27632 on the surface of the Matrigel after it has been cured. (B) The 3D structure of the BBB-in-a-CUBE was visualized by labelling astrocytes with GFP, pericytes with RFP, and BMECs with CFP, and imaging was taken (i) of the 3D view, (ii) at the level of the BMEC sheet, (iii) at the level just below the BMEC sheet, and (iv) in the Matrigel region. On Day 2 after seeding, the astrocytes and pericytes were mostly still rounded, but from Day 4 and Day 6, they can be seen to elongate and populate the region beneath the BMEC sheet, indicating self-organisation of the BBB structure.

Although primary or immortalized cell lines of animal origin were commonly used in many previous BBB models due to their easier availability, high TEER and low permeability values compared to human cells ^43^, iPSC-derived cells, particularly iPSC-derived BMECs based on the work by Lippmann *et al*. ^7,31^, are currently the preferred cell source due to their essentially unlimited supply obtainable by simple differentiation protocols and negation of human-animal species differences, whilst still having high TEER and low permeability. With differentiation protocols for iPSC-derived astrocytes and pericytes also increasingly being developed ^44–48^, fully iPSC-derived models, particularly patient-derived models, could be developed to study human diseases *in vitro* and provide for personalized medicine in the future.

Matrigel was selected as the hydrogel scaffold for the cellular components of the BBB to be grown on as it contains many of the basement membrane components of native BBB such as collagen IV, laminin, nidogen, and perlecan ^49,50^. However, the actual composition of Matrigel is not very well-defined, and there are often reported batch-to-batch inconsistencies with Matrigel supply ^51^. Additionally, though the thickness of the *in vivo* basement membrane is about 30∼100 nm ^50,52^, the thickness of the Matrigel in the BBB-in-a-CUBE is about 1.6 mm, necessitated because of the volume required for the gel to be able to adhere on the CUBE frame by surface tension. However, based on the Day 2 to Day 6 imaging evidence of astrocytes and pericytes migrating towards the surface of the Matrigel closer to the BMECs, it appears that even in a thick Matrigel, the cells can self-organise to form the actual BBB at the surface of the gel. Nevertheless, with the biomaterials field continually evolving to develop alternative biomimetic materials, it may be prudent to consider alternatives such as chemically defined synthetic hydrogels ^51^ or an ultra-thin membrane with ECM hydrogel recently developed ^53^.

### Tight Junctions of BBB-in-a-CUBE

The tight junctions of the BBB act as a physical barrier in blocking the passive diffusion of substances from the blood to the brain ^54,55^. Immunofluorescence staining show that the BBB-in-a-CUBE expresses claudin-5 (CLDN5) and zonula occludens-1 (ZO-1) (Fig. 3A), and qPCR confirmed the mRNA expressions of CLDN5, ZO-1, occludin (OCLN), and junctional adhesion molecule (JAM-A) (Fig. 3B).

**Figure 3.**
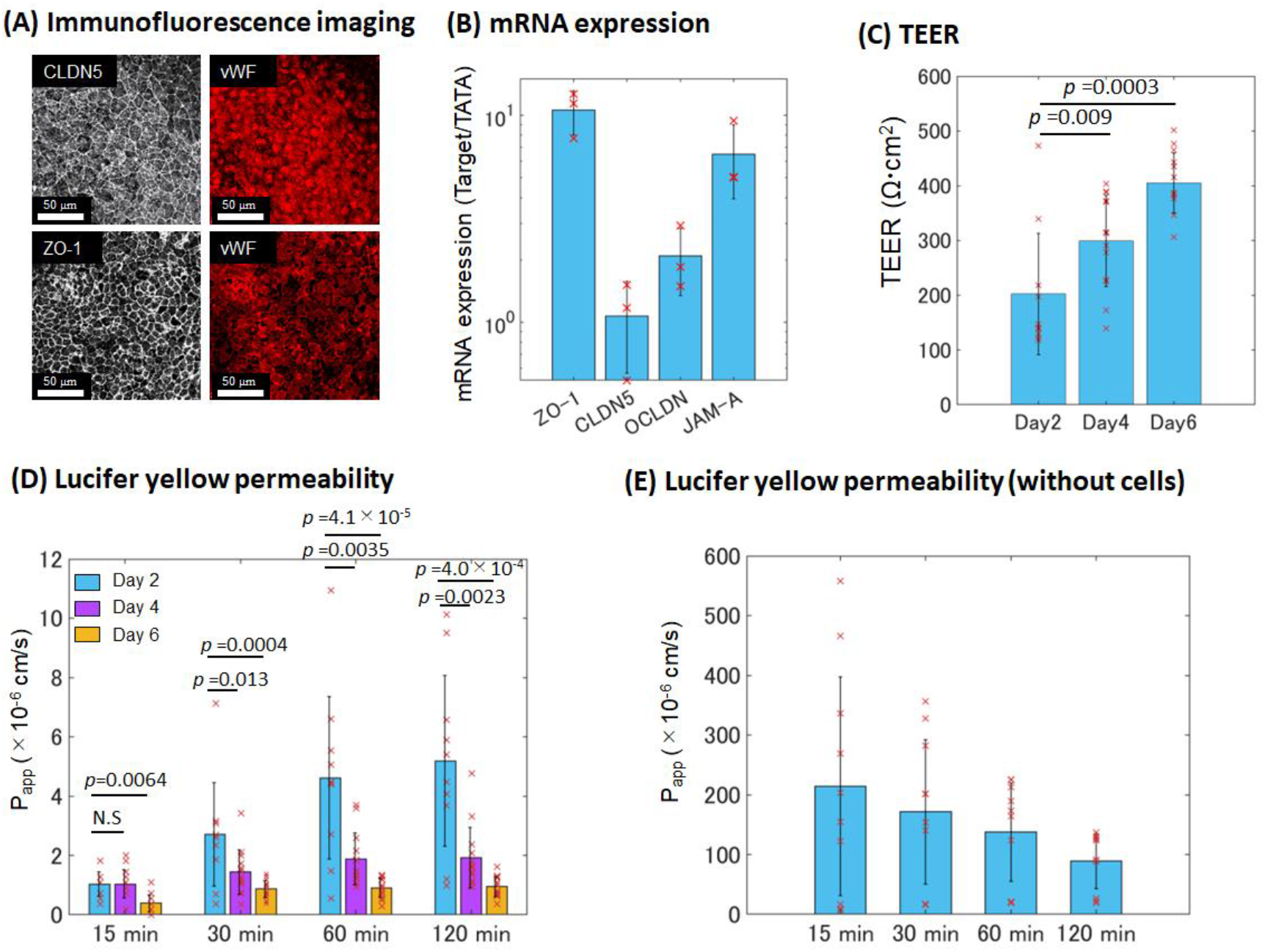
Tight junctions of BBB-in-a-CUBE. (A) Immunofluorescence staining of the tight junction proteins claudin 5 (CLDN) and zonula occludens-1 (ZO-1) with von Willebrand Factor (vWF) marking the BMECs. Scalebar = 50 μm. (B) mRNA expression levels of ZO-1, CLDN5, occludin (OCLN), and junctional adhesion molecule (JAM-A) by RT-qPCR. Expression levels were normalized to that of the house-keeping gene TATA binding protein. (C) Measurement of trans-endothelial electrical resistance (TEER) as a representation of the strength of BBB tight junction showed increasing TEER values over 6 days of BBB-in-a-CUBE culture, indicating stronger barrier formation. (D) The apparent permeability (Papp) of the BBB-in-a-CUBE to small hydrophilic molecule Lucifer Yellow (LY) decreased over the course of 6 days of culture, confirming the strengthening of tight junction function. (E) The much higher Papp values obtained when the LY permeability experiment was performed using Matrigel without any cells confirmed that the low Papp values were not due to accumulation of LY in Matrigel. Data was collected from 3 biological replicates with 6 samples for each replicate for TEER and LY permeability with BBB-in-a-CUBE, and from 1 replicate with 10 samples for LY permeability without cells. p value was calculated by Kolmogorov-Smirnov (KS) test.

TEER measurement is a commonly used method to assess the integrity of endothelial barrier to passive ionic diffusion ^43^. The TEER values measured for BBB-in-a-CUBE were 202 ± 110 Ω.cm^2^ on Day 2, 299 ± 84 Ω.cm^2^ on Day 4, and 405 ± 55 Ω.cm^2^ on Day 6 (Fig. 3C), showing increasing TEER as the cells are cultured for longer. These results are in contrast to other BBB models using iPSC-derived BMEC where TEER values over 1000 Ω.cm^2^ are typically reported, but they tend to peak at about day 1 or 2 before gradually decreasing ^7,31,53,56,57^. A possible reason for this difference is the substrate on which the cells are seeded. While TEER calculations are based on cell seeding area, in the other studies, BMECs were seeded on a porous synthetic membrane, where only 10-15% of the seeding area is actually permeable to ions. On the other hand, BMECs in BBB-in-a-CUBE were seeded on Matrigel only where the whole surface area is permeable to ion movement. The higher resistance of synthetic membranes to electrical current may thus explain the higher TEER values compared to hydrogel only substrate, and it has been reported that BMECs cultured on collagen gels in Transwells (∼840 Ω.cm^2^) showed lower TEER than those cultured directly on Transwells (∼5500 Ω.cm^2^) ^58^. On the other hand, the contribution of astrocytes and pericytes to the formation and maintenance of a tight barrier may explain the increasing TEER in BBB-in-a-CUBE. Although astrocytes and pericytes have been included in several BBB models and resulted in tighter barrier function, the astrocytes and pericytes were often in separate compartments of the Transwell and not interacting directly with one another or with the basement membrane ^5,7,31^. In the BBB-in-a-CUBE model, the close interaction of the three cell types with the basement membrane may have allowed the cells to mature and self-organise to form a more stable BBB, and this may be an interesting avenue to pursue in future work investigating the potential synergistic effects of BBB components in health or disruptive effects in disease.

The movement of small hydrophilic molecules such as Lucifer Yellow (LY) across the BBB is also often used as an indicator of barrier integrity to passive diffusion of non-electrolytes ^43,59^. The LY apparent permeability (P_app_) of BBB-in-a-CUBE was 1∼5 × 10^−6^ cm/s at Day 2, 1∼2 × 10^−6^ cm/s at Day 4, and < 1 × 10^−6^ cm/s at Day 6 (Fig. 3D). Inversely correlating to the increasing TEER from day 2 to day 6 of culture, the P_app_ of BBB-in-a-CUBE to LY decreases and also shows lower variability with longer culture, indicating strengthening of barrier function. As barrier permeability of about 1 × 10^−6^ cm/s are typically used for drug permeability assessments ^56,60^, BBB-in-a-CUBE at Day 6 were used for subsequent studies. To confirm that the low permeability results were not due to accumulation of LY in the Matrigel, LY permeability in Matrigel only without cells was also assessed and showed much higher permeability than BBB-in-a-CUBE samples (Fig. 3E). The decreasing LY permeability from 15 min to 120 min in samples without cells may be due to the negatively charged LY becoming trapped in the gel ^61,62^, but the amount of LY build-up required for this effect is much higher than the amount that passes through the BBB, so we can consider that the low permeability of BBB-in-a-CUBE was a result of BBB function and not because of LY accumulation in Matrigel.

### Transporters of BBB-in-a-CUBE

The transporters of the BBB mediate the movement of essential nutrients from the blood to the brain, as well as the efflux of toxic or unwanted substances from the brain into the bloodstream to be eliminated ^54,55^. Immunofluorescence imaging with scanning from the top BMEC side shows that the BBB-in-a-CUBE expresses multidrug resistance protein 1, also known as p-glycoprotein (MDR1/PGP/ABCB1), multidrug resistance-associated protein 1 (MRP1/ABCC1), breast cancer resistance protein (BCRP/ABCG2), glucose transporter 1 (GLUT1/SLC2A1), large amino acid transporter 1 (LAT1/SLC7A5), monocarboxylate transporter (MCT1/SLC16A1), organic anion transporter (OAT3/SLC22A8), and serotonin (SERT) (Fig. 4A), and qPCR confirmed the mRNA expressions of MDR1, MRP1, BCRP, GLUT1, LAT1, MCT1, OAT3, and aquaporin 4 (AQP4) (Fig. 4B). The localization of transporters in a cell is also important for their proper functioning. To examine the localization of transporters in BBB-in-a-CUBE, the sample was removed from the CUBE frame and cryo-sectioned in the direction perpendicular to the BMEC layer to image the cells from the side view. Immunofluorescence staining results of the sectioned samples show that PGP, BCRP, MRP1, and OAT3, which are efflux transporters, had higher expressions on the luminal (blood) side, while GLUT1, MCT1, SERT, and LAT1 were expressed on both luminal and abluminal (brain) sides of the BMECs (supplementary figure S2), these results are in agreement with previously reported studies ^63,64^.

**Figure 4.**
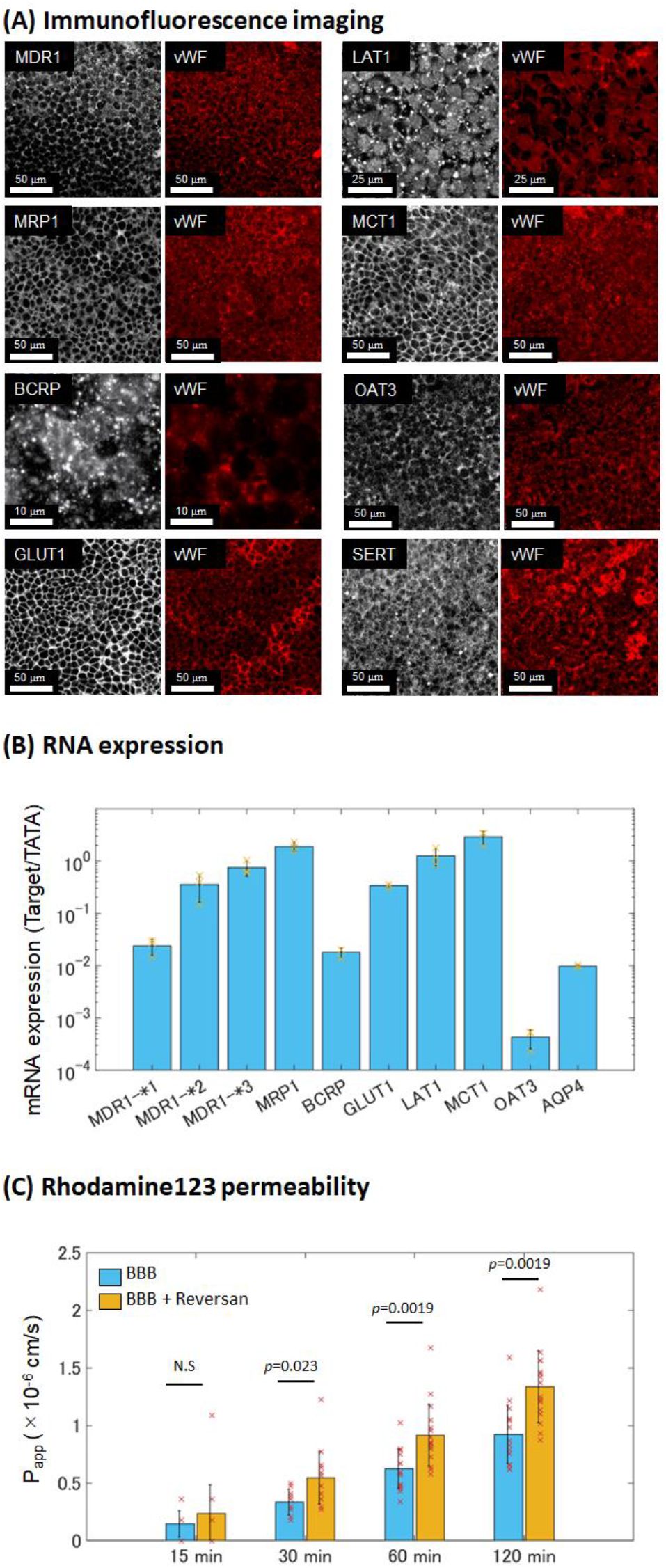
Transporters of BBB-in-a-CUBE. (A) Immunofluorescence staining of the transporters multidrug resistance protein 1 (MDR1/PGP/ABCB1), multidrug resistance-associated protein 1 (MRP1/ABCC1), breast cancer resistance protein (BCRP/ABCG2), glucose transporter 1 (GLUT1/SLC2A1), large amino acid transporter 1 (LAT1/SLC7A5), monocarboxylate transporter (MCT1/SLC16A1), organic anion transporter (OAT3/SLC22A8), and serotonin (SERT) with von Willebrand Factor (vWF) marking the BMECs. Scalebar = 10 μm for BCRP, 25 μm for LAT1, and 50 μm for the others. (B) RT-qPCR confirmed the mRNA expressions of MDR1, MRP1, BCRP, GLUT1, LAT1, MCT1, OAT3, and aquaporin 4 (AQP4). Expression levels were normalized to that of the house-keeping gene TATA binding protein. (C) The apparent permeability (Papp) of the BBB-in-a-CUBE to the PGP substrate Rhodamine123 (Rho123) increased with the addition of the PGP inhibitor Reversan, indicating that BBB-in-a-CUBE expresses functioning PGP transporters. Data was collected from 3 biological replicates with 6 samples for each replicate for Rho123 permeability with BBB-in-a-CUBE. p value was calculated by Kolmogorov-Smirnov (KS) test.

The permeability of Rhodamine123 (Rho123), a PGP substrate, across the BBB barrier is oftentimes used as an indicator of PGP transporter function ^5,7,53,65^. The common method is to measure the efflux ratio of Rho123, which is the ratio of basal-apical flux to apical-basal flux. However, due to the thickness of the Matrigel and potential accumulation of Rho123 in the Matrigel, the efflux ratio may not be representative of the PGP transporter function. Instead, we used the method comparing the permeability of Rho123 with and without the PGP inhibitor Reversan. The Rho123 P_app_ of BBB-in-a-CUBE increased with the addition of Reversan (Fig. 4C), indicating that the presence and function of PGP. It should be noted, however, that Reversan is also an inhibitor for MRP1 ^29,66^ and Rho123 has been reported to be a substrate for organic anion transporting polypeptide (OATP) and organic cation transporters (OCT) transporters ^67,68^. Nevertheless, the significant difference between P_app_ of Rho123 with and without Reversan show that the BBB-in-a-CUBE possesses functioning transporters. More specific investigations beyond the scope of this study, given the large number of transporters that exist, would be required to ascertain whether certain transporters are expressed and functioning in BBB-in-a-CUBE.

### BBB-Glioblastoma Interaction

To demonstrate the application of BBB-in-a-CUBE in drug testing, we designed a chip to contain a BBB-in-a-CUBE and a Glioblastoma-in-a-CUBE as a brain cancer model to test the permeability and effect of the chemotherapy drug Vincristine, based on the study of Tivnan *et al*. ^29^ that reported that inhibition of PGP and MRP1 transporters by Reversan increased the effect of vincristine, a PGP and MRP1 substrate ^5,7,63^, on glioblastoma cell death. At the appropriate timing to start the drug testing experiment, the two CUBEs were assembled side-by-side in the chip, and vincristine drug added to the BBB side of the assembly (Fig. 5A). After 4 hrs, the Glioblastoma-in-a-CUBE was retrieved from the chip and cell death confirmed by live/dead staining. Without the presence of BBB-in-a-CUBE, the percentage of dead T98G glioblastoma cells in Glioblastoma-in-a-CUBE was 9.6%, whereas with BBB-in-a-CUBE, only 6.1% of cells were dead. When Reversan was added with vincristine, the percentage of dead increased to 8.9% (Fig. 5B). These results were in agreement with the report that the inhibition of PGP and MRP1 transport allowed vincristine to pass through the BBB and into the brain where it could exert its effect on the brain cancer. Additionally, the multi-tissue platform developed here requires no pump systems or external power source, thus eliminating the complicated setup often associated with microfluidic devices.

**Figure 5.**
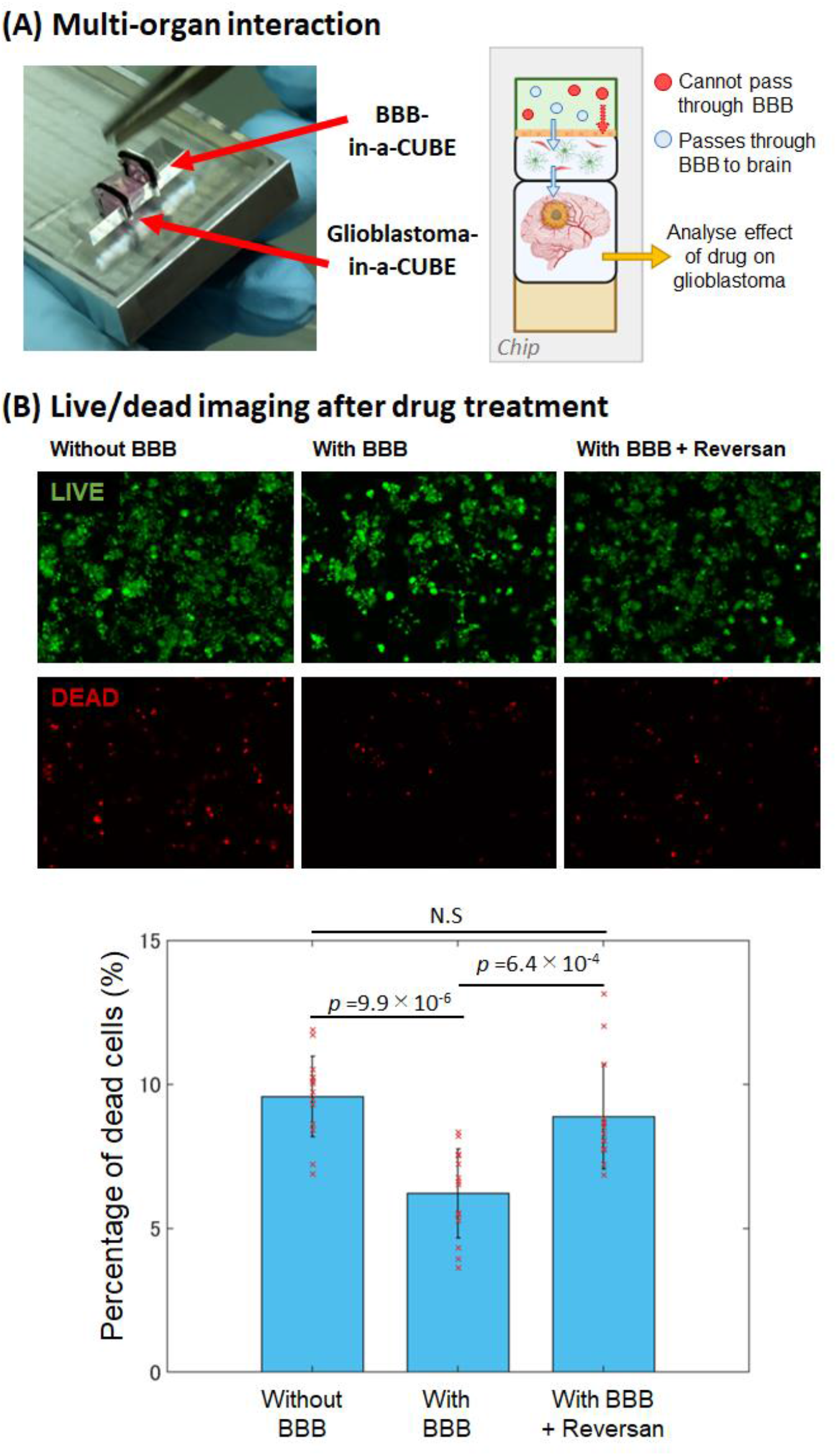
BBB-Glioblastoma interaction. (A) Conceptual diagram of how multi-organ interaction can be achieved by integrating Tissue-in-a-CUBEs together in a chip device. A BBB-in-a-CUBE and a Glioblastoma-in-a-CUBE are positioned together in a chip with their respective growth media in the respective media chambers. Drugs to be tested for treating glioblastoma are then added to the BBB side to determine if they can pass through the BBB, and their effects on the glioblastoma. (B) The proof-of-concept was performed using T98G glioblastoma cells with vincristine, a PGP substrate chemotherapy drug used to treat glioblastoma, as the test drug. The percentage of dead cells was higher when there was no BBB and when the PGP transporter was inhibited with Reversan, compared to when a BBB was present, showing that the drug does not pass through the BBB easily but can be permitted through when the transporter function is inhibited. The results also demonstrated that the effects of the drug on the glioblastoma can be determined by retrieving and analysing the Glioblastoma-in-a-CUBE post experiment. Data was collected from 3 biological replicates with 4∼5 samples for each replicate for live/dead imaging in each condition. p value was calculated by Kolmogorov-Smirnov (KS) test.

Besides its potential application in testing the barrier permeability of a drug and its effect on the target organ, we also envision usage of this platform to study the interaction of different tissues in healthy and diseased states. For example, proper functioning of the BBB is necessary to maintain a healthy brain by regulating homeostasis and clearing substances harmful to the brain from the brain. However, when the BBB becomes disrupted in diseases such as in neurodegenerative diseases, these harmful substances build up in the brain and further disrupts the BBB, causing the disease to worsen progressively ^3,15,69^. By culturing the BBB together with a brain organoid developed from iPSCs of a diseased patient in this multi-organ platform, it would be possible to investigate the interplay between the diseased brain and BBB, and vice versa.

The advantage of this platform is that the tissue samples can be cultured separately as individual components before being integrated into a chip device for experiments; this is particularly relevant for tissues that require long culture times. The differentiation of stem cells to organoids typically requires weeks to months to culture for the organoids form, and not all organoids mature successfully. By culturing organoids in the CUBE device, only organoids that are deemed to have matured successfully can be selected to be incorporated in the multi-organ chip at the appropriate timing, thereby reducing wastage compared to if the organoid had been cultured in the chip from the beginning. Furthermore, the modularity of the tissue samples means that they can be easily retrieved later on for further experiments or post-experiment analyses, increasing the usability of each sample.

One of the disadvantages of this platform in its current form is the low throughput due to the design of the chip - the commercially available clamp holder required to make the device water-tight and prevent leakage around the CUBE during drug testing could only allow us to fit two sets of test chambers in the device. However, we anticipate increasing the number of test chambers with future customisation and improvement of clamp holder and chip designs to enable higher throughput.

Despite the many advantages of PDMS including biocompatibility, permeability to most gases, optical clarity, and simple fabrication method, the use of hydrophobic PDMS as a material component of the CUBE and chip may also be a cause of concern as proteins and small molecules have been known to be absorbed on the surface of PDMS ^70–72^. Nevertheless, several methods have been reported to overcome this issue, including adding poly(ethylene glycol) (PEG) to PDMS to increase its hydrophilicity ^73^ or applying Teflon, 2-methacryloyloxyethyl phosphorylcholine (MPC) polymer or paraffin wax as a coating ^74–76^. Furthermore, the CUBE and chip can also be fabricated using acrylic, which although reduces slightly the optical clarity of the devices, does not absorb proteins on its surface.

### Concluding Remarks

In this study, we established an Organ-on-a-Chip platform utilizing our previously developed CUBE culture device to culture modular tissues or organs that can be assembled in a chip to recapitulate multi-organ interactions. Compared to traditional Transwell systems, the CUBE and chip platform enables the reconstruction of complex 3D tissues or organoids in the CUBE device and the integration of multiple CUBEs to achieve even higher complexity. It is anticipated that the platform can be widely adopted by biology-based researchers and further developed into complex *in vitro* model systems that can reduce the reliance of animal models in both basic research and drug testing.

## Author Contributions

I.K. and M.H. conceived the study, designed and conducted experiments, and analysed the results. I.K. wrote the manuscript. All authors read and approved the submitted manuscript.

## Acknowledgements

This research was supported by funds from JST START Program grant JPMJST1911, and JSPS KAKENHI grants 21H01299, 21K18048. Some illustrations were generated using Biorender.

## Data Availability

The data that support the findings of this study are available from the corresponding author upon reasonable request. Source data underlying graphs in Figures 3B, 3C, 3D, 4B, 4C and 5B are provided in Supplementary Data 1.

---

**Supplementary Figure S1.**
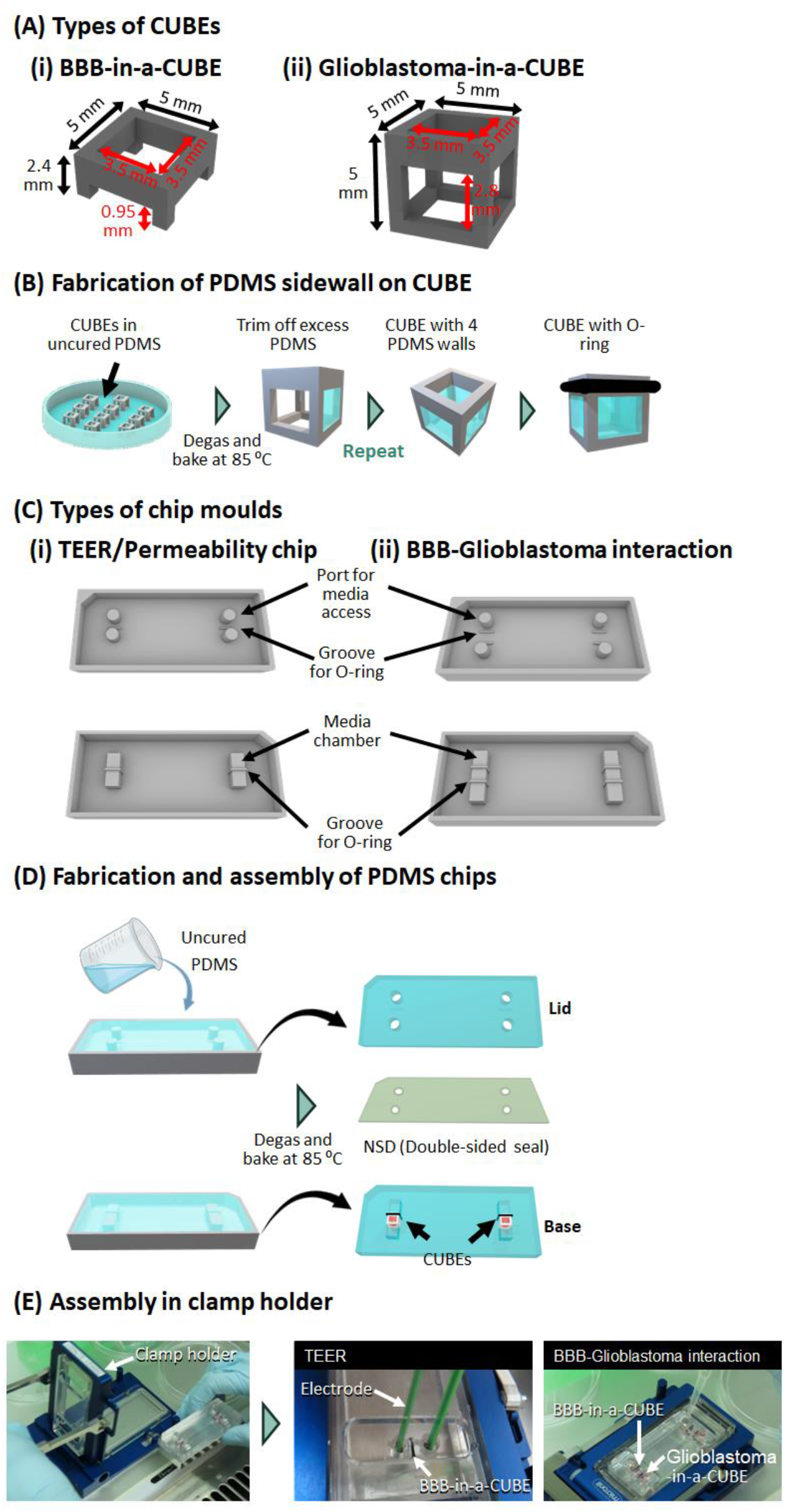
Schematic diagram of fabrication and methods processes. (A) Two different types of CUBEs were used in this study: (i) BBB-in-a-CUBE and (ii) Glioblastoma-in-a-CUBE with different dimensions to suit the different types of tissues being reconstructed. (B) Process to adhere PDMS sidewalls to CUBE device. (C) Two different types of chips were used in this study: (i) chip for TEER and permeability measurements and (ii) chip for BBB-Glioblastoma interaction. (D) Process to fabricate and assemble the PDMS chips. (E) Method to assemble the PDMS chips in a clamp holder for TEER or BBB-Glioblastoma interaction.

**Supplementary Figure S2.**
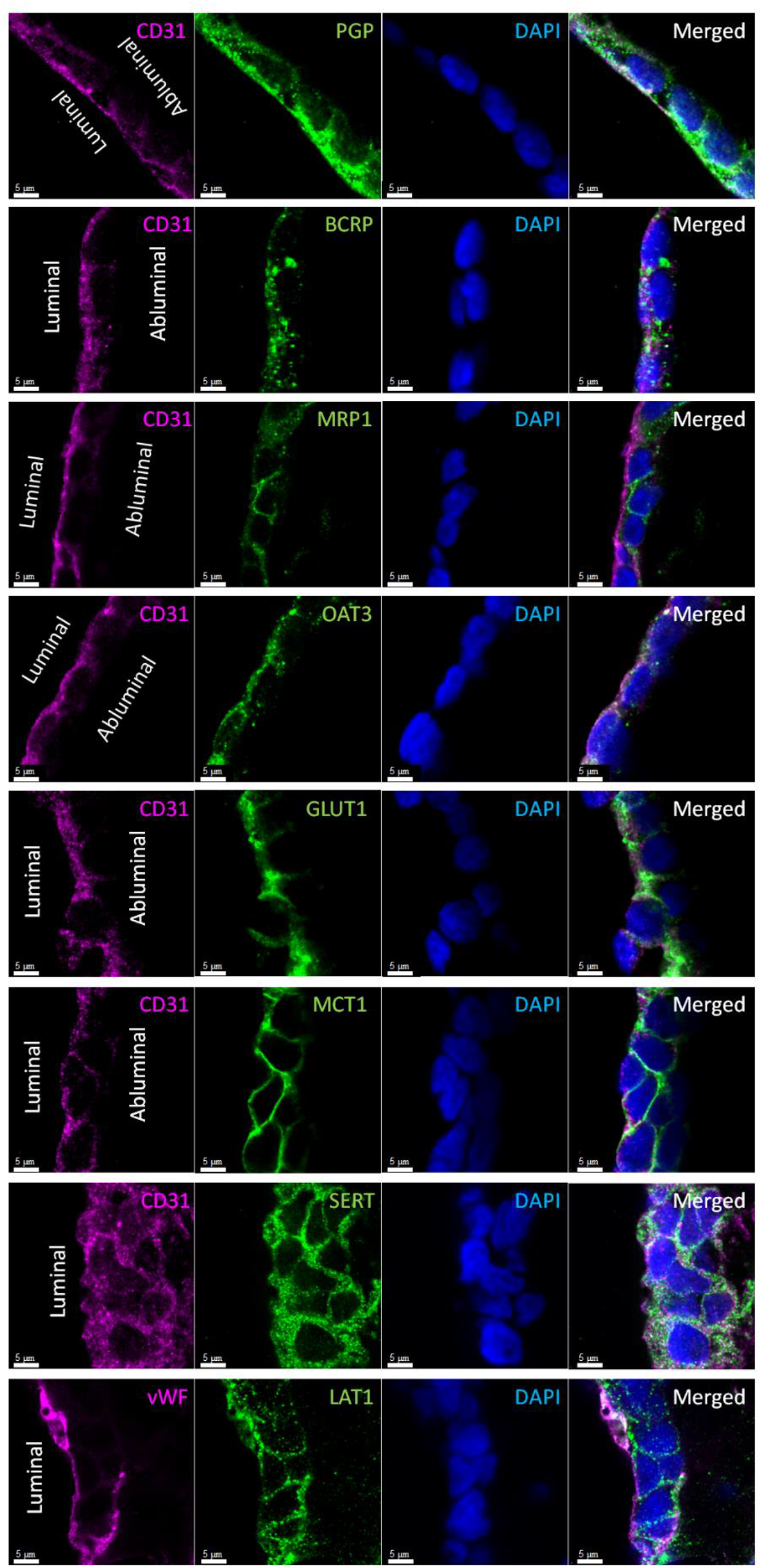
Localization of transporter proteins. Immunofluorescence staining of BMECs from the side view of cryo-sectioned samples show PGP, BCRP, and MRP1, and OAT3 had higher expressions on the luminal (blood) side, while GLUT1, MCT1, SERT, and LAT1 were expressed on both luminal and abluminal (brain) sides of the BMECs. Scalebar = 5 μm.

Supplementary video SV1: Experimental procedure of Modular-Tissue-in-a-CUBE platform.

Supplementary video SV2: 3D imaging of BBB in a CUBE after day 2. BMEC in Blue (CFP), astrocyte in green (GFP), and pericyte in red (RFP) fluorescent respectively.

Supplementary video SV3: 3D imaging of BBB in a CUBE after day 4. BMEC in Blue (CFP), astrocyte in green (GFP), and pericyte in red (RFP) fluorescent respectively.

Supplementary video SV4: 3D imaging of BBB in a CUBE after day 6. BMEC in Blue (CFP), astrocyte in green (GFP), and pericyte in red (RFP) fluorescent respectively.

